# The RNA polymerase clamp interconverts dynamically among three states and is stabilized in a partly closed state by ppGpp

**DOI:** 10.1101/278838

**Authors:** Diego Duchi, Abhishek Mazumder, Anssi M. Malinen, Richard H. Ebright, Achillefs N. Kapanidis

## Abstract

RNA polymerase (RNAP) contains a mobile structural module, the “clamp,” that forms one wall of the RNAP active-center cleft and that has been linked to crucial aspects of the transcription cycle, including loading of promoter DNA into the RNAP active-center cleft, unwinding of promoter DNA, transcription elongation complex stability, transcription pausing, and transcription termination. Crystal structures and single-molecule FRET studies establish that the clamp can adopt open and closed conformational states; however, the occurrence, pathway, and kinetics of transitions between clamp states have been unclear. Using single-molecule FRET (smFRET) on surface-immobilized RNAP molecules, we show that the clamp in RNAP holoenzyme exists in three distinct conformational states: the previously defined open state, the previously defined closed state, and a previously undefined partly closed state. smFRET time-traces show dynamic transitions between open, partly closed, and closed states on the 0.1-1 second time-scale. Similar analyses of transcription initiation complexes confirm that the RNAP clamp is closed in the catalytically competent transcription initiation complex and in initial transcribing complexes (RP_ITC_), including paused initial transcribing complexes, and show that, in these complexes, in contrast to in RNAP holoenzyme, the clamp does not interconvert between the closed state and other states. The stringent-response alarmone ppGpp selectively stabilizes the partly-closed-clamp state, inhibiting interconversion between the partly closed state and the open state. The methods of this report should allow elucidation of clamp conformation and dynamics during all phases of transcription.

**SIGNIFICANCE STATEMENT:** The clamp forms a pincer of the RNA polymerase “crab-claw” structure, and adopts many conformations with poorly understood function and dynamics. By measuring distances within single surface-attached molecules, we observe directly the motions of the clamp and show that it adopts an open, a closed, and a partly closed state; the last state is stabilized by a sensor of bacterial starvation, linking the clamp conformation to the mechanisms used by bacteria to counteract stress. We also show that the clamp remains closed in many transcription steps, as well as in the presence of a specific antibiotic. Our approach can monitor clamp motions throughout transcription and offers insight on how antibiotics can stop pathogens by blocking their RNA polymerase movements.

## INTRODUCTION

RNA polymerase (RNAP) is the main molecular machine responsible for transcription. In bacteria, RNAP is a multi-subunit protein with an overall shape that resembles a crab claw. Comparison of high-resolution structures of RNAP in different crystal lattices (1-5) reveals that one of the two “pincers” of the RNAP crab claw, the “clamp”, can adopt different orientations relative to the rest of RNAP, due to up to ∼20° swinging movements of the clamp about a molecular hinge at its base, the RNAP “switch region” (5-9). The observation of multiple RNAP clamp conformations suggested that the RNAP clamp is a mobile structural module and raised the possibility that RNAP clamp movements may be functionally important for transcription (5-9). In particular, it has been proposed that the RNAP clamp opening may be important for loading DNA into the RNAP active-center cleft and that RNAP clamp closing may be important for retaining DNA inside the RNAP active-center cleft and providing high transcription-complex stability in late stages of transcription initiation and in transcription elongation (5-12).

Single-molecule FRET (smFRET) studies assessing RNAP clamp conformation in solution in freely diffusing single molecules of *Escherichia coli* RNAP confirmed that the RNAP clamp adopts different conformational states in solution; defined equilibrium population distributions of RNAP clamp states in RNAP core enzyme, RNAP holoenzyme, transcription initiation complexes, and transcription elongation complexes; and demonstrated effects of RNAP inhibitors that interact with the RNAP switch region on RNAP clamp conformation (9). However, because those smFRET studies analyzed freely diffusing single molecules--for which it is difficult to monitor individual single molecules over timescales of >1 ms, and thus it has not been possible to define trajectories of smFRET vs. time over timescales of >1 ms--these smFRET studies provided no information about the occurrence, pathway, and kinetics of interconversions between clamp conformational states.

An smFRET study assessing RNAP clamp conformation in surface-immobilized molecules of RNAP from the hyperthermophilic archaeon *Methanocaldococcus jannaschii* showed both open and closed clamp conformations (13). However, interconversions between open and closed clamp dynamics were not observed, possibly due to “freezing” of conformational states at the temperature at which smFRET analysis was performed, a temperature ∼65^°^C below the temperature optimum for the hyperthermophilic archaeal RNAP studied.

Here, we report an smFRET analysis of the RNAP clamp comformation in immobilized single molecules of *E. coli* RNAP at 22°C, a temperature at which *E. coli* RNAP exhibits high activity, under conditions that define trajectories of smFRET vs. time over timescales of up to ∼20 s. We resolve three clamp conformational states, including a previously unresolved partly closed state; we define pathways and kinetics of inter-conversion between clamp conformational states; we confirm clamp equilibrium population distrubutions and define clamp conformational dynamics in transcription initiation complexes; and we show that the clamp conformation can be rapidly and quantitatively remodelled by small-molecule inhibitors and effectors, including the stringent-response alarmone ppGpp. Our work advances our mechanistic understanding of the role of the RNAP clamp in transcription and sets the stage for real-time monitoring of RNAP clamp conformation and conformational dynamics in all phases of transcription.

## RESULTS

### Single-molecule FRET analysis of RNAP clamp conformation and dynamics

To monitor the conformation of clamp in real-time, we have developed a method of surface-immobilizing doubly labeled RNAP molecules (14, 15), and observing the RNAP structure by smFRET using a total-internal reflection fluorescence (TIRF) microscope equipped with alternating-laser excitation (ALEX) (14, 16, 17).

To enable FRET monitoring, the RNAP holoenzyme was labelled with a donor and acceptor at the tips of the β’ clamp and β pincer, respectively, following procedures similar to (18); briefly, fluorophores were site-specifically attached to 4-azidophenylalanine residues introduced to the β and β’ subunits using unnatural amino-acid mutagenesis, followed by Staudinger ligation yielding a fluorescently labelled hexahistidine-tagged RNAP-σ^70^ holoenzyme. This labelling scheme corresponds to a distance of ∼73 Å between the sites of incorporation considering structures of the *E. coli* RNAP holoenzyme (4YG2; Fig. 1A); since this distance is near the Förster radius of the donor-acceptor pair used (∼60 Å; (9)), and that since clamp closing or opening is expected to change the donor-acceptor distance by 5-10 Å, we reasoned that we should be able to use smFRET to study clamp conformational changes in real-time, as we were able to do for the opening-closing motions of the fingers subdomain of bacterial DNA polymerase I (19).

**Figure 1.**
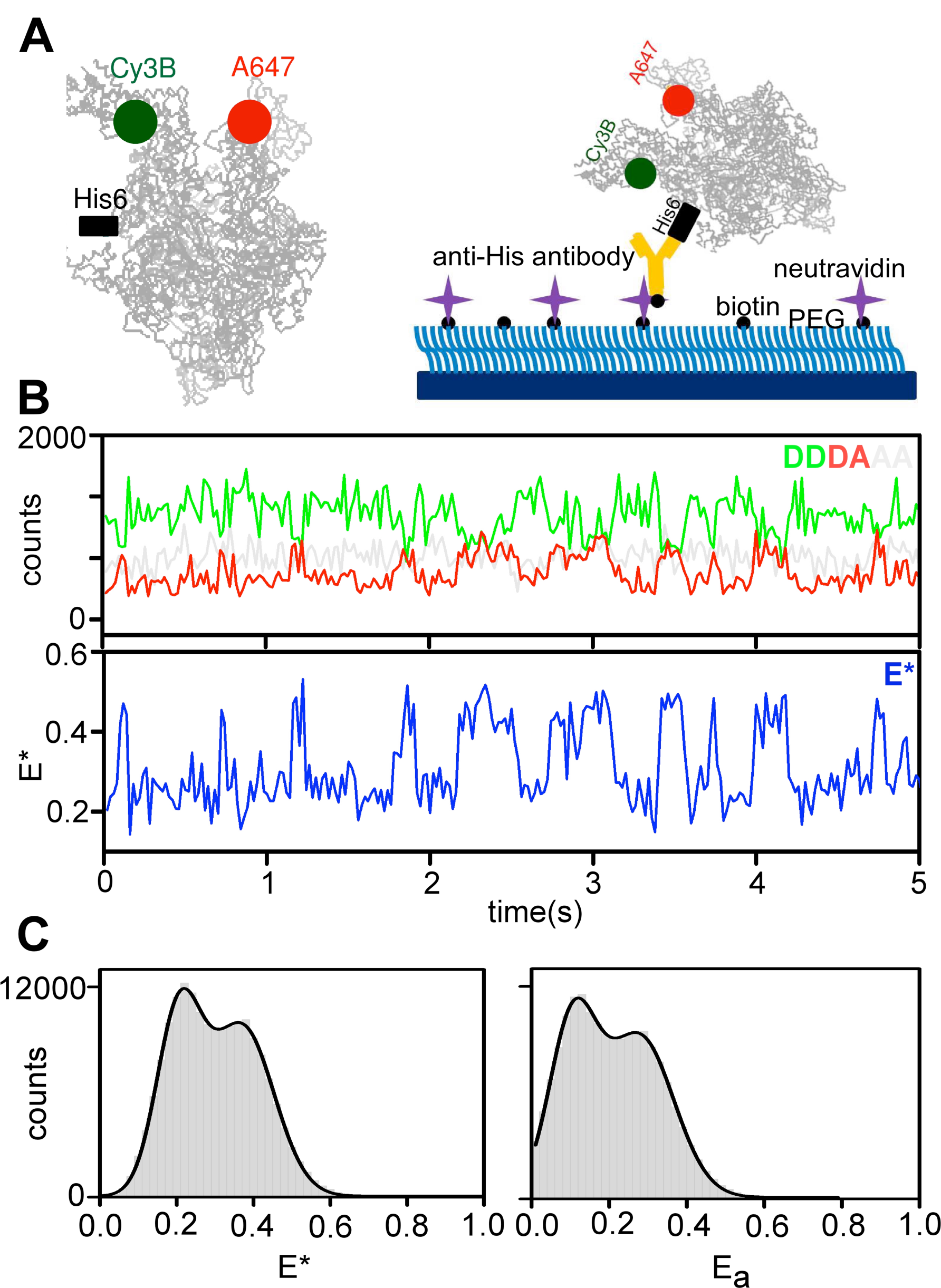
A single-molecule FRET strategy to detect real-time conformational changes of the RNAP clamp on surface-immobilized molecules (A) Left panel: labeling strategy used for detecting clamp conformation changes. Positions of the FRET donor (Cy3B) and acceptor (Alexa647), as well as the position of the β’ C-terminal hexahistidine tag are shown; the rest of the protein (based on the 4YG2 RNAP structure) is shown in grey. Right panel: schematic of the antibody immobilization strategy of RNAP holoenzyme. (B) Example time-trace for an RNAP molecule exhibiting dynamic behavior. The top panels show the fluorescence intensity of the donor emission upon donor excitation (DD; green line) and the acceptor emission upon donor excitation (DA; red line); the clear anticorrelation of the DD and DA signals are consistent with FRET fluctuations involving a single FRET pair. The bottom panel shows the E* trace (in blue) calculated from the intensities. Frame exposure time: 20 ms. (C) RNAP holoenzyme E* histogram showing a bimodal distribution of E* values (left), and accurate E values (right). Data acquisition were performed in T8 imaging buffer at 22°C.

To surface-immobilize molecules of the labelled RNAP, we attached them to PEG-passivated slides coated with antibodies targeting the C-terminal hexahistidine-tag on the β’ subunit of RNAP (Fig. 1B; see (14)). This immobilization strategy preserves the structural integrity and transcription activity of the immobilized RNAP molecules (14, 15, 20). Control experiments also showed minimal non-specific RNAP adsorption to a surface lacking the penta-His antibody (Fig. S1A-B).

### Clamp conformation and dynamics in RNAP holoenzyme: conformation

Following RNAP immobilization, we performed TIRF measurements at 22 ° C with a 20-ms temporal resolution, and used the donor and acceptor fluorescence signals to calculate FRET efficiencies. An initial visual inspection revealed molecules showing stable FRET signals, and molecules fluctuating between different FRET states (Fig. 1B). We selected molecules (*N*=602; each lasting for >50 frames, i.e., >1 s) for further analysis based on robust selection criteria that eliminate aggregates of molecules, as well as molecules with complex photophysics (see *Methods*), and generated an uncorrected FRET (E*) histogram (Fig. 1C, left). The FRET histogram shows an apparent bimodal distribution, with peaks at E* ∼ 0.21 and E* ∼ 0.37; after corrections for cross-talk and detection-correction biases, we also obtained the distribution of *corrected* FRET efficiencies E_a_ (Fig. 1C, right), where the two peaks are found at E_a_ ∼ 0.12 and E_a_ ∼ 0.27.

Our surface-based TIRF microscopy results were similar to previous smFRET studies of diffusing RNAP molecules showing the presence of open and closed clamp conformations (9), with two main differences. First, the solution-based FRET study had also reported a structural state with FRET values higher than what was expected for the closed clamp state; this FRET state was attributed to a collapsed clamp state (with E_a_ ∼0.4). However, we do not see evidence for a significant fraction of a collapsed state in our surface-based studies. We examined possible explanations for this discrepancy, and found that generating a FRET histogram from RNAP holoenzyme molecules *prior* to molecule selection yields a distribution that resembles the solution-based FRET distribution (Fig. S1C). A closer inspection of individual time-traces showed that many molecules in the TIRF-based FRET histogram display behaviours incompatible with single RNAP molecules with a donor and acceptor probe (i.e., correspond to RNAP aggregates or molecules which exhibit complex photophysics; Fig. S1D). Since studies of diffusing molecules do not permit the type of robust filtering of fluorescent species possible in TIRF studies, the confocal-based FRET distributions may have included RNAP aggregates that gave rise to the high-FRET population attributed to the collapsed state.

Second, in addition to molecules showing stable values of E* ∼ 0.2 (Fig. 2A, left) and E* ∼ 0.4 (Fig. 2A, right), we observed many molecules exhibiting a stable intermediate-FRET value with E* ∼ 0.3 (Fig. 2A, middle). Additionally, many molecules were dynamic, featuring clear interconversions between different FRET states (Fig. 2B-C, panels) and anti-correlated changes in their donor and acceptor signals (Fig. 1C). Visual inspection of the time-traces revealed that many interconversions involved the E* ∼ 0.3 FRET state, and one or both of the other FRET states. These results strongly suggest the presence of three clamp conformational states that are able to interconvert to one another in the context of RNAP holoenzyme.

**Figure 2.**
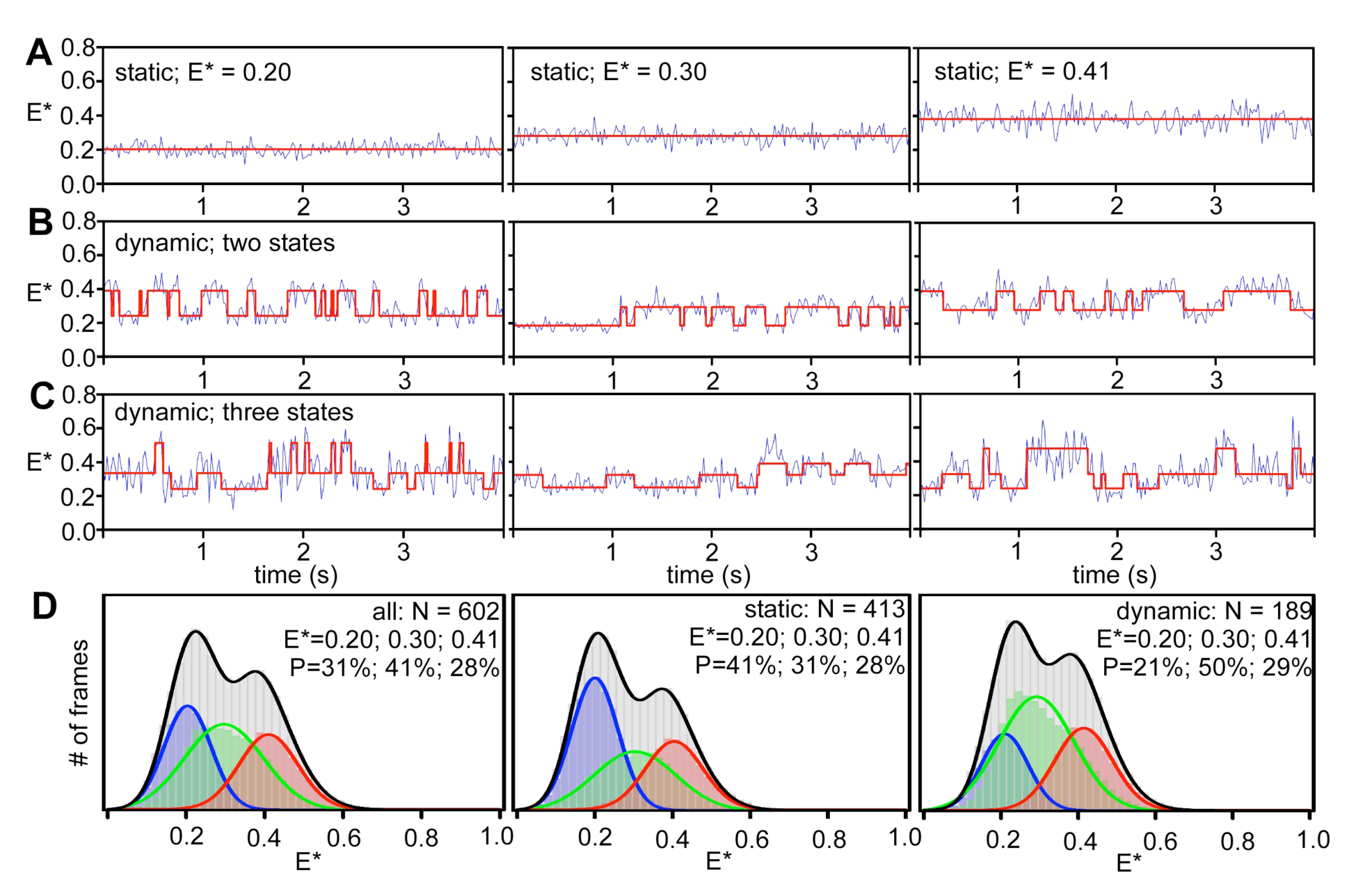
Clamp conformational dynamics in RNAP holoenzyme. Representative time-traces of E* of individual RNAP molecules showing: (A) a stable FRET state at ∼0.2 (fully open clamp; *top left*), a stable FRET state at ∼0.3 (partly open clamp; *middle left*), and stable FRET at ∼0.4 (closed clamp; *bottom left*); (B) transitions between two clamp conformational states; and (C) transition between all three clamp conformational states. In all the panels, E* traces are depicted in blue, and the Hidden Markov Model fits to the data are depicted with red lines. Frame exposure time: 20 ms. (D) E* distribution histogram for a three-state HMM analysis showing relative abundances of fully open clamp (blue), partly open clamp (green) and closed clamp (red) conformations for all molecules (left), for stable molecules only (middle) and for dynamic molecules only (right). Data acquisition was performed in T8 imaging buffer at 22°C.

### Clamp conformation and dynamics in RNAP holoenzyme: dynamics

To analyze further the conformational dynamics of the clamp, we applied Hidden Markov Modeling (HMM) to our time-traces using ebFRET (21); this analysis compares idealised time-traces (assuming a number of states and transition rates between states) with experimental data to produce a lower bound (L) for the log evidence, a quantity that identifies the best among candidate models (21). To identify the number of FRET states that best describe our experimental data, we fit them to models featuring two, three, four, five, or six states (Fig. S2A), and extracted the mean L values for each model; L reached a higher value for the three-state model than the two-state one (Fig. S2B, top), and did not increase significantly upon further increase in the number of states. To compare the models quantitatively, we used the Aikake information criterion (AIC; (22)) which “penalizes” fits with increasing number of states; a plot of AIC versus the number of states also supported the three-state model (a lower AIC value indicates a better fit of the model to the data; Fig. S2B, bottom). We conclude that our FRET results are best described by a three-state model (Fig. 2D, left).

The three-state HMM analysis identified FRET states centered on E* ∼ 0.2, E* ∼ 0.3, and E* ∼ 0.41 (Fig. 2D, left, and showed that ∼68% of the RNAP molecules were static (i.e., they appear to occupy a single FRET state for the 20-s of observation), and ∼32% were dynamic. The FRET states distributed differently between dynamic and static molecules (Fig. 2D, middle and right panels), with the dynamic molecules showing a higher occupancy of the E* ∼ 0.3 state. Dynamic molecules showed interconversions either between two states (22%; Fig. 2B) or amongst all three states (10%; Fig. 2C).

Using the traces of the dynamic molecules, we extracted the dwell-time distributions of the clamp in the three states, fitted them to exponential functions, calculated the average dwell time for each state (Fig. S2C). The transition rates for the interconversion between the three clamp conformational states were extracted from HMM analysis for the three-state model (Fig. S3B). All three dwell-time distributions fitted well to single-exponential functions, and all had lifetimes in the 150-350 ms range. Direct transitions between all three FRET states were observed at similar rates (Fig. S3A, left). The overall RNAP conformational landscape and transition rates (Fig. S3A-B) were largely independent of the way RNAP was reconstituted (i.e., *in vitro* reconstitution vs. *in vivo* reconstitution) and the buffer conditions used (Fig. S3A-B).

### Clamp conformation and dynamics in RNAP holoenzyme: assignment of FRET states to structural states

To assign identified FRET states to RNAP clamp structural states, we compared the observed FRET efficiencies (E_a_), and calculated FRET efficiencies for a series of modelled RNAP clamp structural states ranging from fully open to closed in 2 ° increrments (procedures and curves relating FRET efficiencies to RNAP clamp structural states in (9)). The results indicate that the E* ∼ 0.2, E* ∼ 0.3, and E* ∼ 0.41 FRET states correspond, respectively, to open, partly closed, and closed clamp states, having clamp rotation angles of 0 °, ∼8 °, and ∼16 ° (Fig. 3).

**Figure 3.**
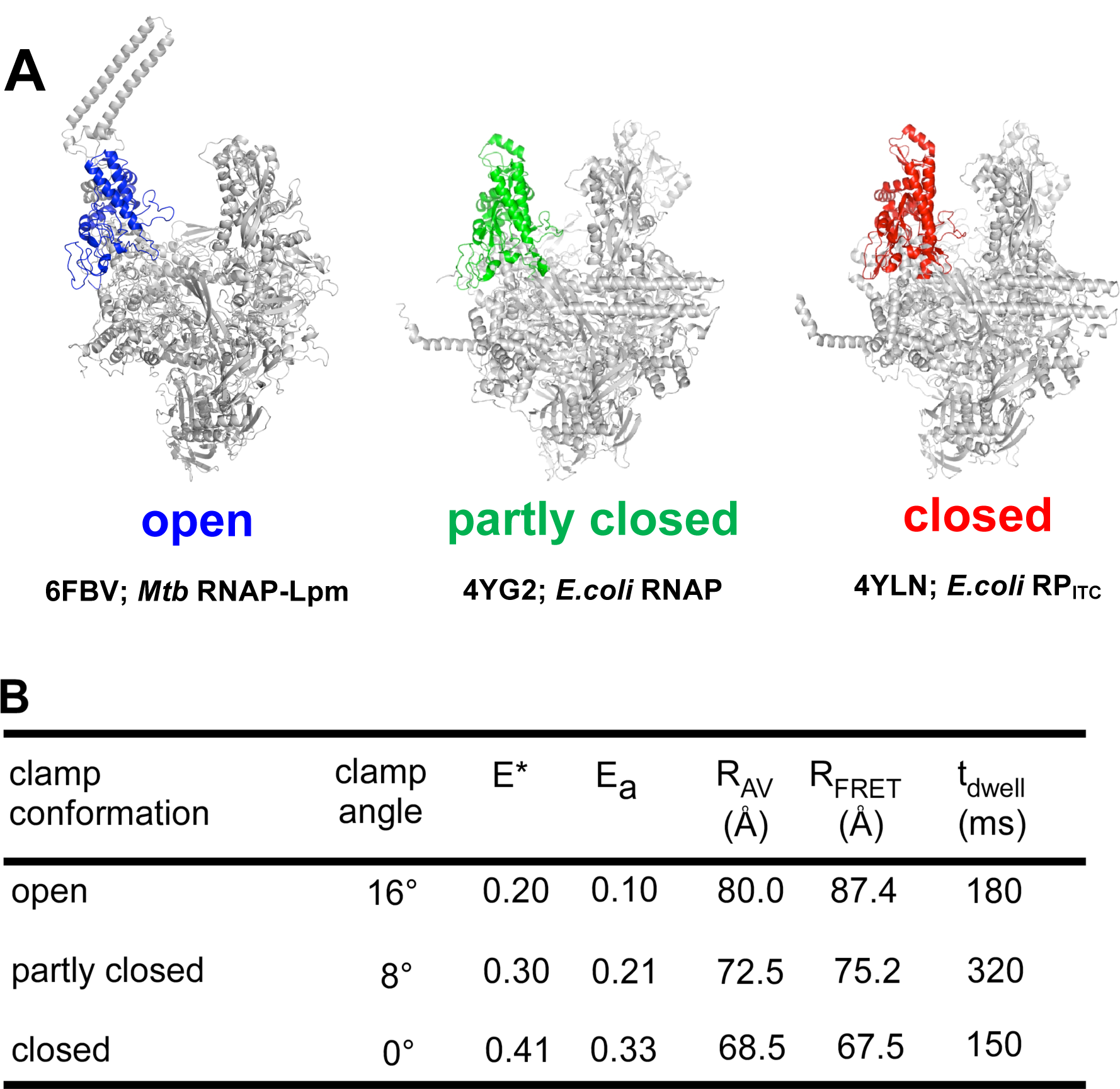
Assignment of FRET states to structural states of RNAP observed in high-resolution structures. RNAP clamp conformation from crystal structures of Mtb RNAP-Lpm complex (left panel; PDB id: 6FBV; RNAP clamp is in blue), RNAP holoenzyme (middle panel; PDB id: 4YG2; RNAP clamp is in green; middle) and RP_ITC_ (right panel; PDB id: 4YLN; RNAP clamp is in red) showing a fully open clamp, a partly closed clamp, and a closed clamp conformation, respectively. (B) Table showing a comparison of clamp angle, measured apparent FRET efficiencies (E*), accurate FRET efficiencies (E_a_), FRET-measured distances (R_FRET_), mean donor-acceptor distances from accessible-volume calculations (R_AV_), and dwell times for the open, partly closed and closed clamp conformations.

The E* ∼ 0.2 FRET state and corresponding open clamp structural state having a clamp rotation angle assigned as 0^°^ match the FRET state observed for *E. coli* RNAP holoenzyme bound to the antibiotic lipiarmycin in solution (Lpm; (5)) and the open clamp structural state observed in a cryo-EM structure of *Mycobacterium tuberculosis* RNAP holoenzyme bound to Lpm (5), as well as in crystal structures of *T. aquaticus* and *T. thermophilus* RNAP core enzymes (PDB 1HQM and PDB 6ASG;(5),(23)).

The E* ∼ 0.3 FRET state and corresponding partly closed clamp structural state having a clamp rotation angle of ∼8 ° match the partly open clamp structural state observed in crystal structures of *E. coli* (PDB 4LK1, 4YG2, 4MEY; (24-26)), and *T. thermophilus* RNAP holoenzymes (PDB 5TMC). This FRET state had not previously been resolved in solution.

The E* ∼ 0.41 FRET state and corresponding closed clamp structural state having a clamp rotation angle of ∼16 ° match the closed clamp structural state observed in crystal structures of *E. coli* (PDB 4YLN; (27)), *M. tuberculosis* (PDB 5UH5; (28)), *M. smegmatis* (PDB 5VI5; (29)), *T. aquaticus* (PDB 4XLN; (30)), and *T. thermophilus* (PDB 4G7H; (31)) transcription initiation complexes and transcription elongation complexes.

### Clamp conformation and dynamics in RP_O._

To assess clamp conformation and dynamics in catalytically competent RNAP-promoter open complexes, RP_O_, we formed RP_O_ at the *lacCONS* promoter (a synthetic consensus σ^70^ bacterial promoter; see *Methods* and Fig. 4A, top), surface-immobilized RP_O_ complexes, and monitored smFRET. The results confirm previous smFRET results indicating that RNAP adopts a closed clamp conformation in RP_O_ ((9); ∼90% occupancy for the E* ∼ 0.41 state; Fig. 4B, top), and show that, within the temporal resolution of our analysis (20 ms), RNAP exhibits static closed-clamp behavior in RP_O_, with no detectable transitions to open or partly closed clamp states (Figs. 4B, S6A).

**Figure 4.**
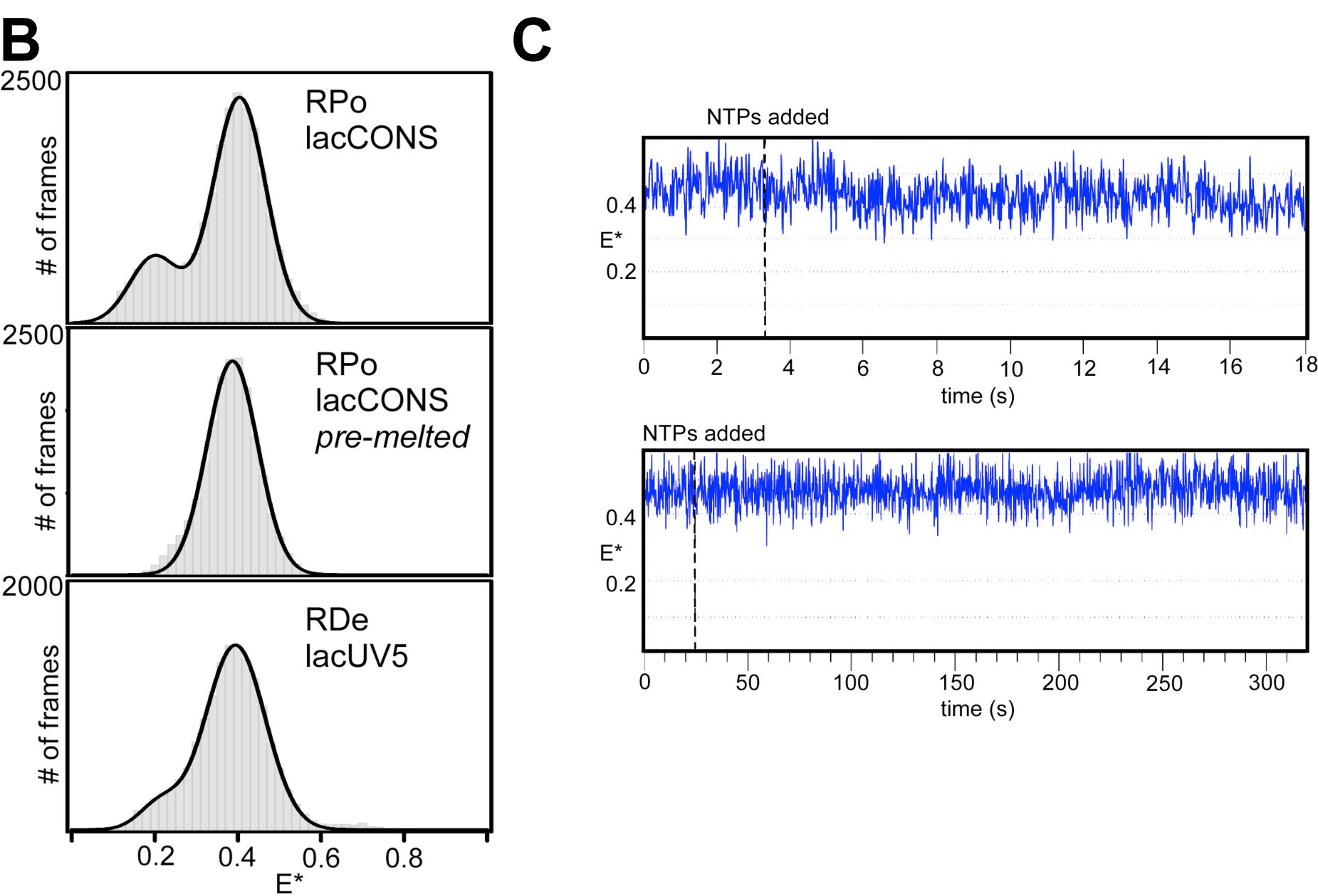
The RNAP clamp conformation remains closed in initial transcription and transcription elongation. (A) Sequences of respective DNA’s are described on the right panels; the -10 and -35 promoter elements are highlighted in dashed boxes, the transcription start sites are highlighted in red and the non-complementary pre-melted DNA sequence in blue. (B) E* histograms (left panels) for RP_O_ formed using a *lac*CONS promoter fragment; for RP_O_ formed using a *lac*CONS promoter fragment having a pre-melted bubble; and for RD_E_ formed on a *lac*CONS-14 promoter fragment using subset of NTPs. Frame exposure time: 20 ms. (C) Representative time traces showing no change in E* values upon addition of ApA and a NTP subset for formation of RP_ITC7_ to RP_O_ formed using a *lac*CONS promoter fragment. Frame exposure times: 20 ms for the top panel, and 200 ms for the bottom panel. For details on the conditions for each complex formation, see *Methods.* Data acquisition was performed in T8 imaging buffer at 22°C.

To assess whether the minor, ∼10% sub-population that adopts an open state indeed corresponds to a non-closed clamp subpopulation of the RP_O_, we performed similar experiments using a “pre-melted” version of the *lac*CONS promoter derivative, in which nontemeplate-DNA-strand and template-DNA-strand sequences at promoter positions -10 to -4 are non-complementary, facilitating RP_O_ formation and increasing RP_O_ stability. This promoter enables analysis of populations in which fully 100% of molecules are in RP_O_, and shows that fully 100% of molecules adopt a closed clamp conformation in RP_O_ (Fig. 4B, middle; Fig. S6B), indicating that the minor, ∼10%, subpopulation of molecules with open clamp states in the experiments in Fig. 4B correspond to molecules that either did not form RP_O_, or that formed RP_O_ and subsequently dissociated.

### Clamp conformation and dynamics in RP_ITC_

To assess clamp conformation and dynamics in RNAP-promoter initial transcribing complexes, RP_ITC_, we surface-immobilized RP_O_ [*lac*CONS promoter (32); see *Methods*)], added an NTP subset that enables RNAP to synthesize RNAs up to 7 nt in length, and monitored smFRET (14, 33). The results confirm previous smFRET results indicating that RNAP adopts a closed-clamp conformation in RP_ITC_ (9) and show that, within the temporal resolution of our analysis (20 ms), RNAP exhibits static closed-clamp behavior in RP_ITC_, with no detectable transitions to open or partly closed clamp states (Figs. 4C, S6C). Since we expect that the majority of RP_ITC_ complexes will be engaged in pausing after the synthesis of a 6-nt RNA during initial transcription (with a pause lifetime of∼15 sec; (14)), and since we do not detect any transitions to open or partly closed clamp states, we conclude that the clamp remains closed during initial transcription pausing.

### Clamp conformation and dynamics in RD_E_

To assess the clamp conformational status and dynamics during transcription elongation, we surface-immobilized RNAP-DNA elongation complexes, RD_E_, [*lac*CONS-14 promoter (32) plus NTP subset that enables synthesis of 14-nt RNA product; see Fig. 4A-B, bottom and *Methods*] and monitored smFRET. The results confirm previous smFRET results indicating that RNAP adopts a closed-clamp conformation in RD_E_ (9), and, as with the result above for RP_O_ and RP_ITC_, show that, within the temporal resolution of our analysis (20 ms), RNAP exhibits static closed-clamp behavior in RD_E_, with no detectable transitions to open or partly closed clamp states (Figs. 4B, bottom; Fig. S6C).

### Effects of small-molecule inhibitors and effectors on clamp conformation: myxopyronin

The clamp conformation is also modulated by the RNAP switch region, which is located at the base of the clamp, serves as the hinge for clamp opening and closing, and forms an important target for RNAP (10, 34-36). To examine effects of a switch-region RNAP inhibitor on clamp conformation and dynamics, we studied effects of the antibiotic myxopyronin (Myx). Myxopyronin binds to the RNAP switch region (9) holoenzyme and inhibits isomerization of the RNAP-promoter closed complex (RP_C_) to RP_O_. Previous smFRET analysis shows that Myx induces or stabilizes a closed clamp conformation (9).

To study effects of Myx on clamp conformation and dynamics, we formed RNAP-Myx complexes in solution, surface-immobilized them, and measured smFRET efficiencies in the absence and presence of Myx. The E* histogram for the complexes showed a single distribution centered at E* ∼ 0.38, corresponding to the closed clamp state, confirming that Myx “locks” the clamp in a closed conformation (Fig. 5C, left). Individual traces of Myx-RNAP complexes did not display any dynamic behavior (Fig. S5A).

**Figure 5.**
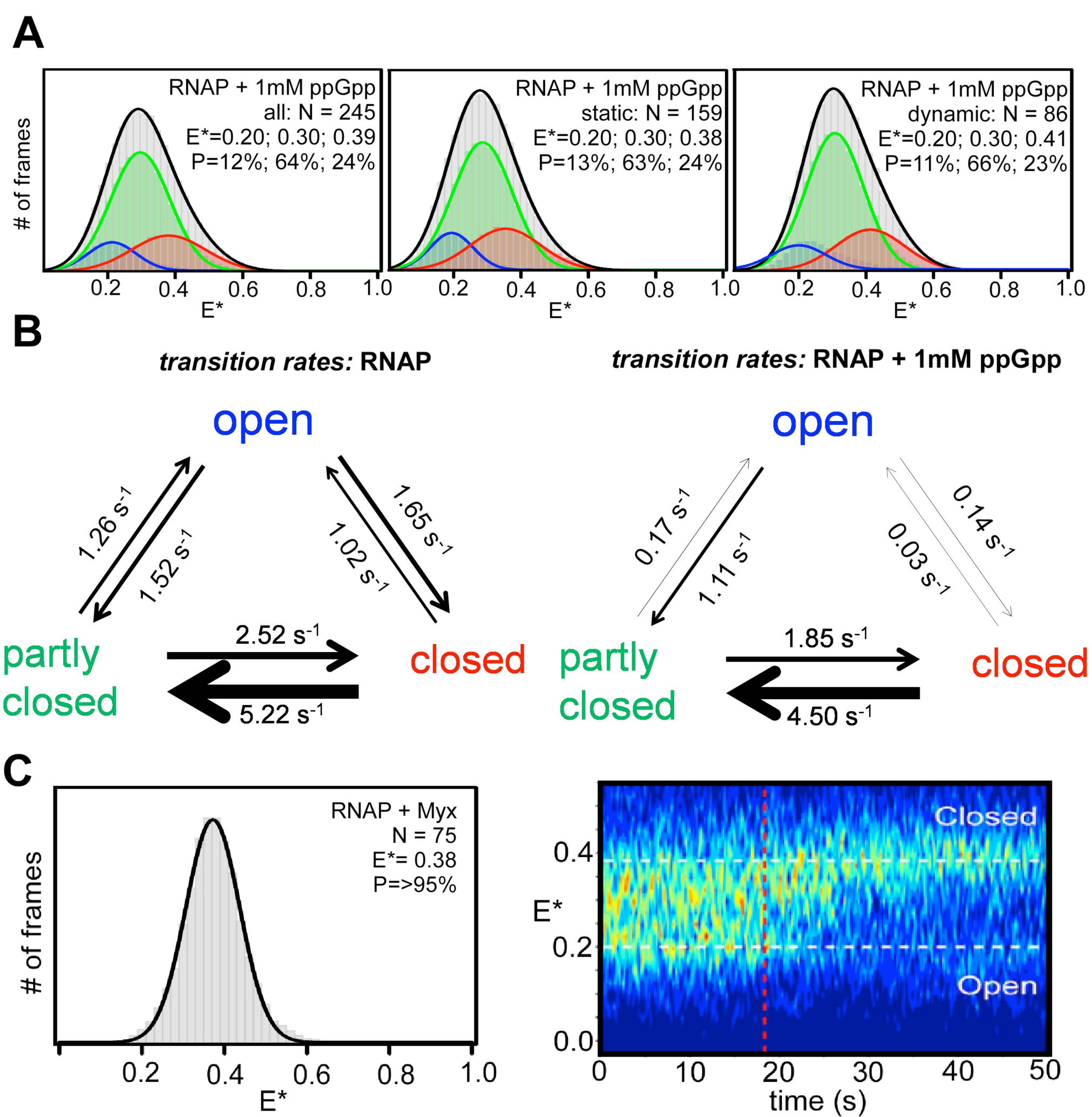
Effect of small molecules on clamp conformation (A) E* histogram for RNAP holoenzyme in presence of 1 mM ppGpp (grey), along with HMM-based E* histograms for the relative abundances of fully open clamp (blue), partly open clamp (green) and closed clamp (red) conformations for all molecules (left), static molecules only (middle) and dynamic molecules only (right). Data acquisition was performed in KG7 imaging buffer at 22°C. (B) Schematic showing rates of transitions between open, partly closed and closed clamp states in absence and presence of 1 mM ppGpp. (C) E* histogram for RNAP holoenzyme in presence of Myx (left) and heat map of E* time traces before and after Myx addition (right). Blue to red colours represent an increasing number of events. Frame exposure times: 20 ms for the left panel, and 200 ms for the right panel. Data acquisition was performed in T8 imaging buffer at 22°C.

To monitor effects of Myx on the clamp conformation and dynamics in real-time, we observed kinetics of FRET changes upon addition of Myx to surface-immobilized RNAP holoenzyme (Fig. 5C, right). Both RNAP molecules initially exhibiting dynamic clamp behavior and RNAP molecules initially adopting static open clamp and partly open conformations transition to a static closed clamp conformation upon addition of Myx (Fig. S5B). The conversion of RNAP holoenzyme molecules to the closed state is relatively slow, taking ∼10 s to convert the initial heterogeneous FRET profile of RNAP to the static closed clamp conformation of RNAP-Myx (see heat map in Fig. 5C, right; Myx added at 18 s; static closed clamp at ∼30 s). We conclude that binding of Myx to RNAP “unlocks” static open and partly open clamp states in favor of a static closed clamp state and substantially reduces the clamp conformational dynamics.

### Effects of small-molecule inhibitors and effectors on clamp conformation: ppGpp

We next examined effects of the stringent-resopnse alarmone ppGpp (37). During the stringent response, ppGpp is produced in high quantities, binds to RNAP, destabilizes RP_O_ at ribosomal RNA (*rrn*) and other promoters, and inhibits isomerization of RP_C_ to RP_O_ at *rrn* and other promoters (38). ppGpp binds to two distinct sites in RNAP, “site 1” located at the interface between RNAP β’ and ω subunits, and “site 2” located near the RNAP secondary-channel β’ rim helices (37, 39-41). It is thought that ppGpp exerts its effects on RNAP through an allosteric mechanism, but details are unclear (37, 40-42).

To assess effects of ppGpp on RNAP clamp conformation and dynamics, we formed RNAP-ppGpp complexes in solution, immobilized them on PEG-passivated surfaces, and measured smFRET efficiencies in the absence or presence of ppGpp. We find that addition of ppGpp reduces clamp conformational heterogeneity, yielding a unimodal FRET distribution centered on E* ∼ 0.3, corresponding to the partly closed clamp state (Fig. 5A, grey distribution). Titration of RNAP with increasing ppGpp concentrations (Fig. S4A) leads to progressive disappearance of open and closed clamp states in favor of the partly closed clamp state. HMM time-traces analysis using a three-state model show greater fractions of molecules in the partly closed clamp state (Fig. 5A, green distribution). For molecules exhibiting dynamic clamp behavior (∼35%, see Fig. S4B), the partly closed clamp state predominates, the closed clamp state is rare, and the open clamp state is very rare (∼66%, ∼23%, and ∼11% of trace durations, respectively; Fig. 5A, right).

Comparison of rates of transition rates between clamp states in the absence and presence of ppGpp shows a large decrease in rates of transitions from partly closed clamp states to the open clamp states (∼8-fold decrease) and in rates of transitions from closed-to-open clamp states and from open-to-closed clamp states (≥30-and ∼12-fold, respectively) upon ppGpp addition. Rates of transitions between partly closed clamp states and closed clamp states do not change. We conclude that ppGpp stabilizes RNAP in the partly closed state and prevents clamp opening. The partly closed state stabilized by ppGpp in solution also corresponds to a state observed in the crystal structures of the RNAP holoenzyme bound to ppGpp (42).

## DISCUSSION

Our analysis of the clamp conformation in the *E. coli* RNAP holoenzyme and its complexes demonstrates the dynamic nature of the clamp and its conformational complexity. Our work extends previous smFRET analysis of freely diffusing single molecules of *E. coli* RNAP, which accessed RNAP clamp conformational landscape in various phases of transcription (9), but which could not observe interconversions due to the short observation time window (∼1 ms). Our study also complements smFRET analysis of RNAP clamp conformation in immobilized single molecules of a hyperthermophilc archaeal RNAP (13), which defined the RNAP clamp conformational landscape, but which did not show dynamic interconversions between clamp states, possibly due to “freezing” of interconversions by performing smFRET analysis at a temperature ∼65°C below the temperature optimum for the hyperthermophilc RNAP studied. Here, we have recorded FRET time trajectories for surface-immobilized molecules of *E. coli* RNAP at temperatures at which *E.* coli RNAP exhibits high activity; we have obtained trajectories of smFRET vs. time on the 20-s timescale; and we have observed and characterized interconversions between clamp states.

We identified considerable heterogeneity among RNAP holoenzyme molecules, which can either be static (stably open, stably partly closed, or stably closed), or dynamic (fluctuating between open, partly open, and closed conformations on the sub-second timescale). Notably, not all molecules are dynamic, despite the long observation spans for each molecule (20 s). We interpret static RNAP states as occupying free-energy local minima separated from other conformational states, including the dynamic state, by significant energy barriers. This pattern of behavior (which has also been seen for static and dynamic transcription-bubble states; (15)) may reflect a slow RNAP conformational change required to “unlock” clamp dynamics).

The results we obtained from dynamic molecules confirmed the long-standing hypothesis that the RNAP clamp is indeed a mobile structural element, switching dynamically between different conformational states. We expect this conformational equilibrium to shift more towards dynamic states, as the temperature increases to 37**°**C, since the conformational changes involved are thermally driven.

We resolved a previously unresolved, partly closed clamp conformational state. The partly closed state we observe in solution corresponds to a state observed in all the crystal structures of the RNAP holoenzyme. We observe that, in solution, ppGpp induces or stabilizes the partly closed clamp conformational state and reduces the clamp-opening rates; we also note that the partly closed state we observe in solution corresponds to a state observed in the crystal structures of the RNAP holoenzyme bound to ppGpp. We infer that ppGpp binds to RNAP in solution and restricts the motions of the clamp in a way that prevents full clamp opening. Since we did not include DksA in the assay, these effects must arise from ppGpp binding to the DksA-independent ppGpp binding site on RNAP (“site 1”) located at the interface between β’ and ω (39, 42).

Our results confirm that, in RP_O_ and RP_ITC_, the clamp is in the closed conformational state, and show that within out temporal resolution (∼20 ms), does not detactably interconvert, even transiently, to open or partly closed states in RP_O_ and RP_ITC_. It should be noted that RP_ITC_ is engaged in iterative cycles of synthesis and release of abortive RNA products (43). Accordingly, the absence of detectable inter-conversion from the closed state to open and partly closed states in RP_ITC_ suggests that the clamp remains stably closed in individual nucleotide-addition cycles and in abortive-RNA release. It also should be noted that, at the promoter analyzed, RP_ITC_ engages in initial-transcription pausing, exhibiting a pause with ∼15 s half life at position +6 (14, 20). Accordingly, the absence of detectable inter-conversion from the closed state to open and partly closed states in RP_ITC_ further suggests that the clamp remains stably closed during initial-transcription pasusing.

The results here and in refs. (5, 9) with the small-molecule inhibitors and effectors Lpm, ppGpp, and Myx provide a set of chemical probes that lock the clamp in, respectively, the open, the partly closed, and the closed clamp conformational states, facilitating analysis of functional properties of these conformational states.

Straightforward extensions of the procedures of this report should enable analysis of effects on clamp conformation and dynamics of other small-molecule inhibitors and effectors, macromolecular regulatory factors, and physiological conditions, and should enable analysis at any stage of transcription, from promoter search through transcription termination.

## MATERIALS AND METHODS

### DNA, RNAP and myxopyronin preparation

Oligonucleotides were purchased from IBA Life Sciences (Germany) and annealed in hybridization buffer (50 mM Tris-HCl pH 8.0, 500 mM NaCl, 1 mM EDTA). Fluorescently labelled, hexahistidine-tagged *E.coli* RNAP holoenzyme (hereafter “labelled RNAP”) with Cy3B and Alexa647 at positions 284 on the β’ subunit, and 106 on the β subunit, respectively, were prepared using *in vitro* reconstitution as described in (9, 18) and *in vivo* reconstitution as described in (5). Myxopyronin was prepared as described in (10).

### Formation of RNAP complexes

RP_O_ was formed as described in (14). Briefly: 250 nM of lacCONS+2 promoter DNA or pre-melted lacCONS+2 promoter (Fig. 5A) was incubated with 50 nM labelled RNAP for 15 min at 37°C in T8 buffer (50 mM Tris-HCl, pH 8.0, 100 mM KCl, 10 mM MgCl_2_, 100 μg/ml BSA, 1 mM DTT and 5% glycerol), followed by addition of 1 mg/ml heparin sepharose, and a further incubation for 1 min at 37 °C. The reaction mixture was centrifuged, the supernatant transferred to pre-warmed tubes at 37°C and further incubated at 37°C for 20 minutes.

RD_E_ was formed as follows: 250 nM of lacCONS-14 promoter DNA fragment (Fig. 5A) was incubated with 50 nM labelled RNAP in T8 buffer supplemented with 0.5 mM ApA, for 15 min at 37 °C, followed by addition of 1 mg/ml heparin sepharose, and further incubation for 1 min at 37 °C. The reaction mixture was centrifuged, the supernatant transferred to pre-warmed tubes at 37 °C and incubated for 20 min at 37°C. A mixture of ATP, UTP and GTP (final concentration of 12.5 μM each) was added to the reaction, and further incubated for 5 min at 37 °C.

### Single-molecule fluorescence instrumentation

Single-molecule FRET experiments were performed on a custom built objective-type total-internal-reflection fluorescence (TIRF) microscope (16). Light from a green laser (532 nm; Samba; Cobolt) and a red laser (635 nm; CUBE 635-30E, Coherent) was combined using a dichroic mirror, coupled into a fiber-optic cable, focused onto the rear focal plane of a 100x oil-immersion objective (numerical aperture 1.4; Olympus), and displaced off the optical axis such that the incident angle at the oil-glass interface exceeds the critical angle, creating an evanescent wave. Alternating-laser excitation (ALEX) was implemented by directly modulating the two lasers using an acousto-optical modulator (1205C, Isomet). Fluorescence emission was collected from the objective, separated from excitation light by a dichroic mirror (545 nm/650 nm, Semrock) and emission filters (545 nm LP, Chroma; and 633/25 nm notch filter, Semrock), focused on a slit to crop the image, and then spectrally separated (using a dichroic mirror; 630 nm DLRP, Omega) into donor and emission channels focused side-by-side onto an electron-multiplying charge-coupled device camera (EMCCD; iXon 897;Andor Technology). A motorized x/y-scanning stage with continuous reflective-interface feedback focus (MS-2000; ASI) was used to control the sample position relative to the objective.

### Single-molecule fluorescence imaging of RNAP and its complexes

For all single-molecule experiments, a biotin-PEG-passivated glass surface was prepared, functionalized with neutravidin and treated with biotinylated anti-hexahistidine monoclonal antibody (Qiagen) as described (14, 15) to form observation chambers with (biotinylated anti-hexahistidine monoclonal antibody)-neutravidin-biotin-PEG/mPEG-functionalized glass floors. For experiments with the RNAP holoenzyme, a 100 pM solution of fluorescently labelled hexahistidine tagged RNAP holoenzyme was incubated on the PEGylated surfaces derivatives with biotinylated anti-hexahistidine monoclonal antibody (15) for 5 min at 22°C in T8buffer; the same protocol was used for surface-immobilizing RP_O_ and RD_E_ complexes. Binding was monitored until ∼30 molecules were deposited on the field of view (10 x 12 μm). Immobilization densities typically were ∼30 molecules per 10 μm x 12 μm field of view. Immobilization specificities typically were >98% (assessed in control experiments omitting biotinylated anti-hexahistidine monoclonal antibody; Fig. S1A-C).

Observation chambers containing immobilised labelled RNAP were washed with 2×30 μl T8, and 30 μl T8 imaging buffer (50 mM Tris-HCl pH 8.0, 100 mM KCl, 10 mM MgCl_2_, 100 mg/ml BSA, 1 mM DTT, 5% glycerol and 2 mM TROLOX, plus an oxygen scavenging system consisting of 1 mg/mL glucose oxidase, 40 μg/mL catalase, and 1.4% w/v D-glucose) was added just before movies were recorded. For experiments monitoring initial transcription in real time, RP_O_ was formed and immobilized as described previously (14) and T8 imaging buffer supplemented with 500 μM ApA was added to the observation chamber just before recording. NTP reaction mixtures (UTP and GTP; final concentrations of 80 μM) in T8 imaging buffer were then added to the observation chamber manually during data acquisition.

For experiments analysing the effects of ppGpp on RNAP holoenzyme, 1 mM ppGpp was added to 30 nM labelled RNAP in KG7 buffer, followed by incubation for 5 min at 37 °C. The sample was then diluted into KG7 buffer supplemented with 1 mM ppGpp, and was added to the surface to surface-immobilize the labelled RNAP in presence of ppGpp at 22°C for 5 min. Observation chambers containing labelled RNAP in presence of ppGpp were washed with 30 μl 2×KG7 buffer supplemented with 1 mM ppGpp, and 30 μl KG7 imaging buffer (40 mM HEPES-NaOH, pH 7.0, 100 mM potassium glutamate, 10 mM MgCl_2_, 1 mM dithiothreitol, 100 μg/ml bovine serum albumin, and 5% glycerol, 2 mM TROLOX, plus an oxygen scavenging system consisting of 1 mg/ml glucose oxidase, 40 μg/ml catalase, 1.4% w/v D-glucose) supplemented with 1 mM ppGpp was added just before movies were recorded. Experiments with RNAP holoenzyme in absence of ppGpp were carried out in KG7 buffer at 22°C in a similar manner for direct comparison of results.

For experiments analysing the effect of Myx on clamp conformation, 20 μM Myx was added to labelled RNAP in T8 buffer, and incubated for 10 min at 37°C. The sample was diluted into T8 imaging buffer supplemented with 20 μM Myx, and incubated on the surface to immobilise the labelled RNAP in presence of Myx at 22°C for 5 min. Observation chambers containing labelled RNAP in presence of Myx were washed with 30 μl 2×T8 buffer supplemented with 20 μM Myx, and 30 μl T8 imaging buffer supplemented with 20 μM Myx was added just before movies were recorded. For real-time Myx binding experiments, labelled RNAP was immobilized as above in T8 imaging buffer, and 20 μM Myx was manually added to the observation chamber during data acquisition. Data acquisition was done at 21°C with exposure times of either 20 ms or 200 ms. For 20-ms temporal resolution experiments, excitation powers of 8 mW at 532 nm, and 4 mW at 635 nm were used. For 200-ms temporal resolution experiments, excitation powers of 0.5 mW at 532 nm, and 0.15 mW at 635 nm were used.

### Image analysis and data processing

Movies of surface-immobilized labeled RNAP were analyzed using the home-built software TwoTone-ALEX (16), and the background-corrected intensity-vs.-time traces for donor emission intensity upon donor excitation (I_DD_), acceptor emission intensity upon donor excitation (I_DA_), and acceptor emission intensity upon acceptor excitation (I_AA_) were extracted as described (16). For each dataset, we manually inspected intensity time traces and exclude traces exhibiting I_DD_ <300 or >2,000 counts or I_AA_ <200 or >2,000 counts; traces exhibiting multiple-step donor or acceptor photobleaching; traces exhibiting donor or acceptor photobleaching in frames 1-50; and traces exhibiting donor or acceptor blinking (Fig. S1D). The set of selected intensity time traces were used to calculate time traces of apparent donor-acceptor FRET efficiency (E*) and donor-acceptor stoichiometry (S), as described (44):

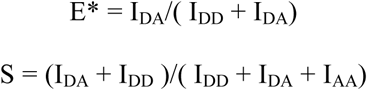

The E* time traces include only data points preceding any donor or acceptor photobleaching events. Two-dimensional E*-S plots were constructed using all selected data points to distinguish species containing donor only (D-only), acceptor only (A-only), and both donor and acceptor (D-A). For species containing both donor and acceptor (D-A), one-dimensional E* histograms were plotted. E* values were corrected, and accurate donor-acceptor efficiencies (E_a_) and donor-acceptor distances (R) were calculated as described (45). Briefly, we correct for the following: leakage (46) of the donor emission into the acceptor-emission channel [calculated from E^*^ values for the donor-only population (E*_D-only_): Lk = E^*^_D-only_/(1-E^*^_D-only_)]; and direct excitation (Dir) of the acceptor by the green laser was calculated from S values of an acceptor only population (S_A-only_): Dir = S_A-only_/ (1-S_A-only_). Correction for the leakage and direct acceptor excitation yields the FRET proximity ratio, E_PR_, which is defined in terms of E*, S, Lk and Dir as follows (16):

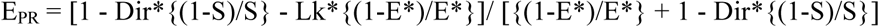

Finally, accurate FRET for each subpopulation is determined from the detection factor, γ (0.95 in this work; determined as γ = ΔI_AA_/ΔI_DD_, where ΔI_AA_ and ΔI_DD_ are changes in I_AA_ and I_DD_ upon acceptor photobleaching) (47) and E_PR_ as follows: E_a_ = E_PR_/ [γ-(γ-1)*E_PR_]. Mean donor-acceptor distances (R) were calculated from mean E values using: R = R_0_ [(1/E_a_)-1]^1/6^, where R_0_ is the Forster parameter (60.1 Å; (9)).

### HMM analysis of FRET time-traces

The set of E* time traces selected for analysis were analyzed using Hidden Markov Modelling as implemented in the ebFRET software ((21); we used 0.2 and 0.4 as min and max prior centers, 0.05 for noise, and 5 iterations), and fitted to models with two, three, four, five, or six distinct E* states (Fig. S2A). Each model was optimized by maximising a lower bound (L) as defined in (21). To select the model that best describes the data, values of lower bound, L, for each model were compared (Fig. S2B). To compare between the models we also estimated the Aikake information criteria (AIC) as described in (22). Both values of lower bound and the AIC vs number of states plot reveal the three state model to be the best fit. Visual inspection of individual time traces and transitions among states also confirmed the presence of three distinct FRET states. The apparent FRET efficiencies (E*) from the fit to a three-state model were extracted, plotted in Origin (Origin Lab) for the overall population, as well as for the static subpopulation alone, and the dynamic subpopulation alone. The resulting E* histograms linked to each state were fitted to single Gaussian functions in Origin. The resulting histograms provide the equilibrium population distributions of states with distinct E*, define numbers of subpopulations with distinct E*, and, for each subpopulation, define mean E* (Fig. 2D).

For dwell-time and transition rate measurements, an HMM analysis was applied to time traces exhibiting dynamic behavior. To identify dynamic traces from the set of selected molecules, individual traces were manually checked and those showing anti-correlated changes in the DD and DA channel were identified. Molecules showing greater than three transitions were considered to be dynamic. The data were fitted to a three-state model, dwell times for each state were extracted and plotted as frequency distributions; the distributions were then fitted to a single-exponential decay in Origin (Fig. S2C), and mean dwell times for the corresponding states were extracted. The rates of transitions between the states were calculated from the transition probabilities (extracted from HMM analysis on ebFRET for the three-state model) between the three clamp conformational states by multiplying the transition probabilities with the number of frames per second (Fig. S3B).

### Accessible-volume modelling and clamp angle calculations

For accessible-volume calculations of the FRET pairs on different RNAP crystal structures, and calculation of predicted distances between donor-acceptor pairs, we used methods described in (46, 48). The position of attachment for Alexa 647 phosphine and Cy3B phosphine were the Cα atom of residue 106 of the RNAP β subunit, and the Cα atom of residue 284 of RNAP β’ subunit, respectively. The Cy3B phosphine dye was characterized by alinker length of 21 Å, a linker width of 4.5 Å, and dye radii of 6.8, 3, and 1.5 Å (*x, y*, and *z*, respectively). The Alexa 647 phosphine dye was characterized by a linker length of 26 Å, a linker width of 4.5 Å, and dye radii of 11, 4.7, and 1.5 Å. The accessible volume of donor and acceptor probes and the average donor acceptor distances for all possible donor-acceptor pair positions were calculated using the FPS software (46). RNAP clamp angles were determined using methods described in (9).

## DATA SHARING STATEMENT

All our fluorescence and FRET time-trace data, as well as the HMM software used for their analysis, will be available to any interested party upon request.

## AUTHOR CONTRIBUTIONS

A.N.K. and R.H.E conceived the project; A.M. prepared and characterized protein samples; D.D., A.M., and A.M.M. performed single-molecule experiments and analyzed data; D.D. and A.M. prepared figures; and D.D., A.M., R.H.E, and A.N.K wrote the manuscript.

## ACKNOWLEDGEMENTS

D.D. was supported by an UK Engineering and Physical Sciences Research Council DTA studentship. A.N.K. was supported by Wellcome Trust (grant 110164/Z/15/Z), and the UK Biotechnology and Biological Sciences Research Council (grants BB/H01795X/1 and BB/J00054X/1). R.H.E. was supported by NIH grant GM041376. A.M.M. was supported by the Instrumentarium Science Foundation, Finnish Cultural Foundation and Alfred Kordelin Foundation.

